# KIS counteracts PTBP2 and regulates alternative exon usage in neurons

**DOI:** 10.1101/2023.05.15.540804

**Authors:** Marcos Moreno-Aguilera, Mònica B. Mendoza, Alba M. Neher, Martin Dodel, Faraz K. Mardakheh, Raúl Ortiz, Carme Gallego

**Affiliations:** Molecular Biology Institute of Barcelona (IBMB), CSIC, Catalonia, 08028, Spain; Centre for Cancer Cell and Molecular Biology, Barts Cancer Institute, Queen Mary University of London, London, UK

**Keywords:** KIS kinase/ PTBP2/ alternative splicing/ exon usage/ neuronal differentiation

## Abstract

Alternative RNA splicing is an essential and dynamic process to control neuronal differentiation and synapse maturation, and dysregulation of this process has been associated with neurodegenerative diseases. Recent studies have revealed the importance of RNA-binding proteins in the regulation of neuronal splicing programs. However, the molecular mechanisms involved in the control of these splicing regulators are still unclear. Here we show that KIS, a brain-enriched kinase with a domain shared by splicing factors, controls exon usage in differentiated neurons at a genome-wide level. KIS phosphorylates the splicing regulator PTBP2 complex and markedly counteracts its role in exon exclusion. At the molecular level, phosphorylation of unstructured domains within PTBP2 causes its dissociation from key co-regulators and hinders its RNA-binding capacity. Taken together, our data provide new insights into the post-translational control of splicing regulators and uncover an essential role of KIS in setting alternative exon usage in neurons.

## Introduction

In the mammalian brain, alternative splicing (AS) is a key mechanism that contributes substantially to the enormous molecular diversity among neuronal cell types (Merkin *et al*, 2012; Barbosa-Morais *et al*, 2012; Mazin *et al*, 2021). Despite the similarity of neuronal gene expression programs, each neuronal subclass can be distinguished by unique alternative mRNA processing events. Moreover, AS is known to drive many aspects of neurogenesis, including neural progenitor cell proliferation and differentiation, axon guidance, synapse formation, and synaptic plasticity (Vuong *et al*, 2016a; Furlanis & Scheiffele, 2018). The relevance of alternative splicing as a regulatory mechanism is evidenced by numerous examples in which alterations in splicing are associated with pathological states (Gandal *et al*, 2018; Schieweck *et al*, 2021; LaForce *et al*, 2022). Although fine-tuning mRNA processing is essential for neuronal activity and maintenance, insights into its regulation are just starting to emerge. RNA-binding proteins (RBPs) play a critical role in regulating alternative splicing patterns. These proteins bind to pre-mRNAs and form ribonucleoprotein (RNP) complexes that can affect splicing by influencing the recognition of splicing signals or by blocking access of the splicing machinery to certain regions of the pre-mRNA. Some RBPs act as splicing enhancers, promoting the inclusion of specific exons, while others act as splicing repressors inhibiting the inclusion of exon inclusion (Grosso *et al*, 2008; Fisher & Feng, 2022; Traunmüller *et al*, 2023).

Among the most investigated RNA-binding proteins are the polypyrimidine tract-binding proteins, PTBP1 and PTBP2, which are considered master regulators of neuronal fate (Keppetipola *et al*, 2012; Linares *et al*, 2015; Zhang *et al*, 2019). These two proteins exhibit tissue-specific expression patterns in which PTBP1 is expressed in most cell types and neuronal progenitor cells, whereas PTBP2 is expressed primarily in differentiating neurons and testis. During the differentiation of neuronal progenitor cells to postmitotic neurons a switch takes place from the predominant expression of PTBP1 to its neuronal paralog PTBP2, which is essential for the stem cell to neuron transition. PTBP2 null mice die shortly after birth and exhibit misregulation of AS in genes involved in cytoskeletal remodeling and cell proliferation (Boutz *et al*, 2007; Licatalosi *et al*, 2012) as well as in neurite growth and synaptic transmission(Li *et al*, 2014). Furthermore, PTBP2 knockout brains display reduced neural progenitor pools and premature neurogenesis (Licatalosi *et al*, 2012). PTBP2 also plays an important role in the embryonic-specific repression of alternative exons. As PTBP2 expression decreases from its highest levels in the embryonic brain to moderate levels during postnatal development, a cohort of exons switches from being skipped in embryos to displaying enhanced inclusion in adults (Licatalosi *et al*, 2012; Zheng *et al*, 2012; Li *et al*, 2014). Thus, sequential downregulation of PTBP1 followed by PTBP2 contributes to the activation of neural exon networks at the appropriate stages of development (Raj & Blencowe, 2015). AS of the *Dlg4* transcript, which encodes the excitatory postsynaptic protein PSD95, is an example of the relevance of the PTBP1/2 switch in neuronal differentiation (Zheng, 2016). Skipping of exon 18 in *Dlg4* transcripts causes premature translation termination and nonsense-mediated mRNA decay (NMD). The controlled induction of this exon during neuronal development is essential to ensure the timing of synapse formation (Zheng *et al*, 2012). This exon may also be dynamically regulated in mature neuronal circuits to adapt PSD95 expression to the requirements of synaptic remodeling (Soto *et al*, 2019).

PTBP1 and PTBP2 are 74% identical in amino acid sequence and have a similar domain organization of four RNA recognition motifs (RRMs) connected by three linker regions. Although primarily characterized as repressive splicing regulators, they can also activate some splice sites as a function of binding position relative to regulated exons (Corrionero & Valcárcel, 2009; Xue *et al*, 2009; Llorian *et al*, 2010). It has been shown that PTBP1/2 RRM2, along with the following linker sequence, is sufficient for splicing repressor activity (Robinson & Smith, 2006). RRM2 can interact with both RNA via its canonical β-sheet surface, and with other co-regulators such as Matrin3, Raver1 and hnRNPM (Coelho *et al*, 2015). Furthermore, Tyr247 of PTBP1 (PTBP2 Tyr244) within RRM2 is particularly critical for co-regulatory protein interactions (Rideau *et al*, 2006; Joshi *et al*, 2011). Regarding the regulation of PTBP proteins, different phosphate modifications are located in the unstructured regions, including the N-terminal, Linker 1 and Linker 2 regions (Pina *et al*, 2018). To date, the kinases involved are still under investigation as well as the role of these post-translational modifications in controlling PTBPs splicing activity.

KIS was first identified as a Kinase that Interacts with Stathmin (Maucuer *et al*, 1997), a phosphoprotein that controls microtubule dynamics (Watabe-Uchida *et al*, 2006). Curiously, KIS is the only known protein kinase that possesses a U2AF homology motif (UHM), an atypical RRM that has lost its RNA-binding ability, but mediates protein–protein interactions at the C-terminal region of KIS (Kielkopf *et al*, 2004; Corsini *et al*, 2007; Manceau *et al*, 2008). This motif shares 42% sequence similarity with U2AF65, a 65 kDa subunit of the splicing factor U2AF, suggesting that KIS could be involved in splicing-related processes. Various lines of evidence support the function of KIS in the modulation of gene expression at different levels such as transcription, splicing, mRNA stability and protein translation (Manceau *et al*, 2008; Pedraza *et al*, 2014; Arfelli *et al*, 2023). Although KIS is ubiquitously expressed, higher levels are detected in the nervous system, and KIS mRNA level increases gradually during postnatal development, reaching its highest level in the mature brain (Bièche *et al*, 2003). In mature neurons, KIS has been implicated in dendritic spine morphogenesis and synaptic activity (Pedraza *et al*, 2014).

Here, we find that KIS kinase is involved in the control of alternative exon usage, a key mechanism for promoting functional gene expression patterns during neuronal differentiation. We show that alternative splicing regulators are the main phosphorylation targets of this UHM-containing kinase. Notably, we find that PTBP2, an essential AS factor for neuronal maturation, is the major phosphotarget of KIS. We further demonstrate that phosphorylation by KIS dissociates PTBP2 protein complexes affecting their ability to bind RNA and therefore inhibiting PTBP2 activity. Interestingly, phosphomimetic mutations are predicted to have a significant impact on the structure of the PTBP2 RRM2 repressor domain. Taken together, our data demonstrate that KIS phosphorylation counteracts PTBP2 activity and thus alters isoform expression patterns of genes involved in neuronal differentiation and synaptic activity.

## Results

### KIS kinase regulates exon usage during neuronal differentiation

KIS contains a UHM domain frequently found in splicing regulatory proteins, and accumulates in nuclear sub-structures adjacent to those formed by splicing factors (Fig 1A). Prompted by its role in proper expression of key postsynaptic proteins (Pedraza *et al*, 2014), we decided to analyze genome-wide exon usage during neuronal differentiation *in vitro*. First, immature neurons from primary cortical neurons of 7 days *in vitro* (DIV) were compared with mature neurons from 18DIV cultures when KIS expression reached maximal levels in hippocampal cultures (Fig 1B). Second, cortical neurons were infected at 7DIV with a lentiviral construct co-expressing GFP and a shRNA targeted against KIS (shKIS) and collected at 18DIV (Fig 1C). Total RNA was purified from triplicate samples for deep sequencing analysis. When compared to control (shCtrl), shKIS-infected cells displayed a five-fold reduction in KIS mRNA levels which led to KIS levels similar to those at onset of neuronal differentiation (Fig 1D). Our analysis revealed that the impact of KIS on exon usage affected both exon inclusion and skipping, although with a clear bias towards exon inclusion 6.3% *vs* 2% skipping. (Fig 1E and Dataset EV1). Notably, genes containing more than one exon with at least a two-fold change coded for proteins significantly enriched in the synapse (Fig EV1A), including two important activators of neurite outgrowth, CAMSAP1, and NCAM1 (Fig EV1B). These data suggest that KIS promotes exon inclusion in genes related to neuronal differentiation, especially genes encoding synaptic proteins. Supporting this notion, the top 100 exons upregulated by KIS, i.e. downregulated by KIS knockdown, were strongly enriched among those increasingly used by cortical neurons from 7DIV to 18DIV (Fig 1F). Among a total of 8457 exons that displayed low variability (Fano factor <5×10^-4^) within the KIS knockdown triplicate samples, 1141 and 1106 exons were found significantly upregulated and downregulated, respectively, by KIS (Fig 1G and Dataset EV2). In addition, 28.0% of exons upregulated by KIS were also upregulated during differentiation, whereas only 18.8% decreased their usage from 7DIV to 18DIV (Fig EV1C). The opposite behavior was observed for exons downregulated by KIS in which 14.3% and 24.8% increased or decreased their usage during differentiation, respectively (Fig EV1C). Finally, whereas genes containing KIS-downregulated exons did not retrieve specific GO terms, exons upregulated by KIS encoded a higher percentage of protein disorder (Fig EV1D) and were found in genes preferentially coding for RNA-binding proteins (Fig 1H), key factors of RNA granule formation and transport along neurites. Overall, our data link KIS to the control of exon usage in cortical neurons and suggest a pivotal role of this UHM-containing kinase as a regulator of alternative splicing during neuronal differentiation.

**Figure 1.**
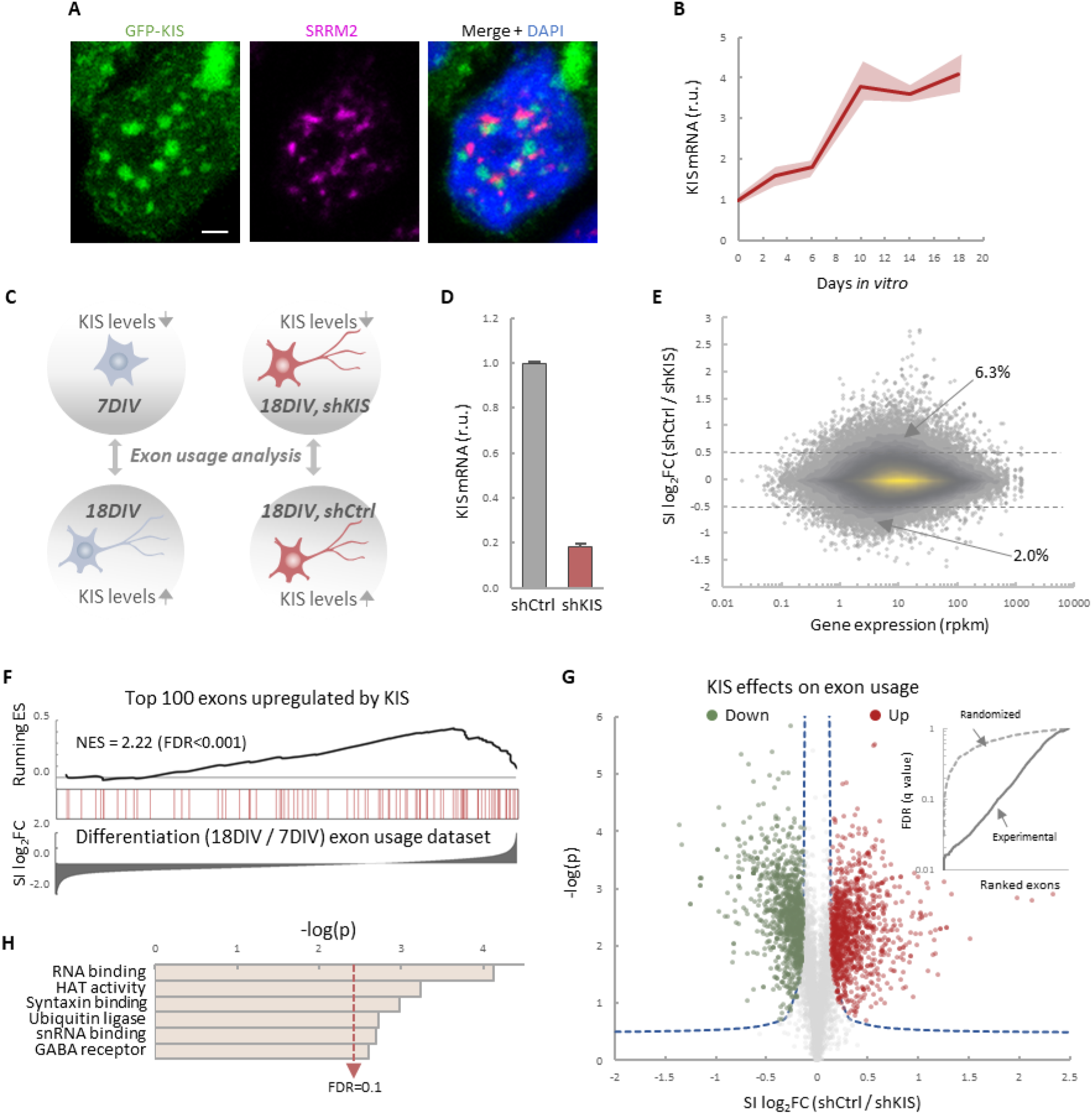
KIS promotes the inclusion of exons upregulated during neuronal differentiation. **A** Immunofluorescence staining of endogenous speckle marker SRRM2 (magenta) and DAPI (bue) in HEK293T cells transfected with GFP-KIS (green). Scale bar, 2 μm. **B** Differential expression of KIS in cultured cortical neurons from E17.5 mouse embryos. Samples were collected at different days *in vitro* (DIV), analyzed by qRT-PCR and made relative to day 0. Mean ± SEM (n = 3) are plotted. **C** Experimental design of exon usage analysis. Cultured cortical neurons from E17.5 mice were collected at 7DIV or 18DIV (left experimental set). Cortical neurons were transduced at 7DIV with lentiviral vectors expressing shCtrl or shKIS and collected at 18DIV (right experimental set). Total RNA was purified from triplicate samples for deep sequencing analysis. **D** KIS mRNA levels in primary cortical neurons transduced at 7DIV with lentiviral vectors expressing shCtrl or shKIS and collected at 18DIV. Bars represent mean ± SEM (n = 3). **E** MA-plot shows splicing index (SI) fold change (log2FC) as a function of gene expression in rpkm (reads per kilobase million) in control neurons (shCtrl) compared to KIS knockdown neurons (shKIS) as in panel d. The percentage of exons displaying a log2FC above 0.5 or below −0.5 is indicated. **F** RNA-seq analysis from cortical neurons expressing shCtrl or shKIS as in panel d and from cortical neurons collected at 7 DIV and 18 DIV for neuronal differentiation (n = 3). Barcode plot shows the position of exons upregulated by KIS, i.e. downregulated by KIS knockdown, within the transcriptomic dataset of exons upregulated during neuronal differentiation. Normalized enrichment scores (NES) and the corresponding FDR values are shown. **G** Volcano plot shows t-test significance (-log2p) as a function of splicing index (SI) fold change (log2FC) in control neurons (shCtrl) compared to KIS knockdown neurons (shKIS). Significantly (FDR < 0.05) downregulated (n = 1106, green) or upregulated (n = 1141, red) exons are highlighted. Inset shows the dependence of FDR on exon rank from experimental and randomized datasets. **H** GO term enrichment analysis of genes with exons upregulated by KIS. FDR value is shown.

### Phosphoproteomics links KIS to RNA splicing and identifies PTBP2 as a novel phosphotarget

One possible way to control the use of alternative exons during neuronal differentiation is by changing the expression of splicing regulators. Our RNA-seq analysis showed that KIS knockdown did not affect transcript levels of splicing regulators whose expression underwent changes during cortical neurons differentiation (Fig EV2A). Therefore, we tested whether the KIS phosphoproteome could reveal the mechanism by which KIS regulates exon usage. For this, we analyzed phosphorylation changes in the proteome of HEK 293T cells transfected with KIS and KIS^K54A^, a point mutant without kinase activity. We chose this cell line because of its high transfection efficiency and, although it is an embryonic kidney-derived line, there is evidence to suggest a neuronal lineage, which could explain the expression of neuron-specific genes (Lin *et al*, 2014). Quantitative phosphoproteomic analysis resulted in the identification of 9994 phospho-sites in a total of 2743 identified proteins (Dataset EV3). As shown in the MA-plot (Fig 2A), proteins displaying the strongest differential phosphorylation by KIS are highly related to alternative splicing, i.e. PTBP2, RAVER1, hnRNPM and MATR3. PTBP2 is considered one of the pivotal splicing regulators in neuronal development (Keppetipola *et al*, 2012) and, strikingly, phosphorylation at serine 308 of PTBP2 is among the top phosphor-sites in our KIS phosphoproteome (log2FC =2.36). Furthermore, the kinase assay confirmed that among the three KIS serine targets on PTBP2 (Ser178, Ser308 and Ser434), Ser308 is key for PTBP2 phosphorylation by KIS (Fig EV2C and D). Ser178 is present in a very large tryptic peptide (37 aa) that was not detected in the KIS phosphoproteome but, as it fulfills the consensus site phosphorylated by KIS (Fig EV2E), we decided to include this residue in our analysis. Note that the interactions between splicing factors that are KIS targets have been already shown except for SUGP1 (Fig 2B and see also Discussion).

**Figure 2.**
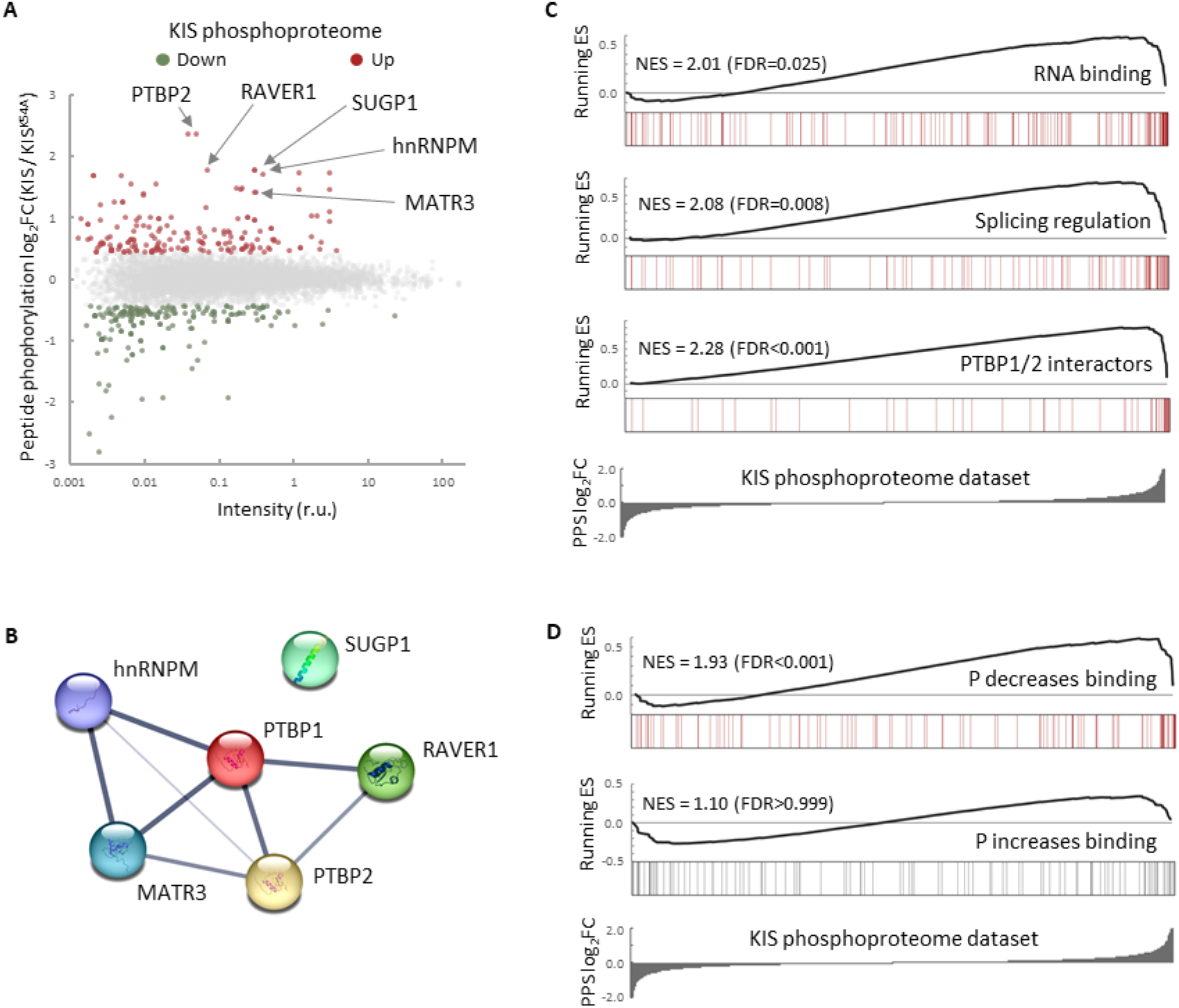
The phosphoproteome of KIS identifies splicing regulators. **A** MA-plot of phospho-sites found in the proteome of HEK293T cells transfected with KIS and KIS^K54A^ mutant. **B** STRING interaction map of top KIS phosphorylated proteins. **C**,**D** Gene enrichment analysis of the KIS phosphoproteomic dataset. Barcode plots indicate the rank position of genes from (C) GO terms and PTBP1/2 interactors and (D) phosphorylated RNA-binding proteins (Vieira-Vieira *et al,* 2022) that were found significantly enriched within the KIS phosphoproteomic dataset. Normalized enrichment scores (NES) and the corresponding FDR values are shown.

Next, to perform Gene Set Enrichment Analysis (GSEA), we ranked KIS targets using a Phospho Protein Score (PPS) that incorporates the fold-change of all identified phosphopeptides per protein (see Materials and Methods). Proteins that are more phosphorylated in the presence of KIS have higher PPS, while those that are more phosphorylated in the presence of KIS^K54A^ have lower PPS. GSEA analysis confirmed that KIS-phosphorylated proteins are enriched in regulation of splicing, RNA binding and PTBP1/2 interactors (Fig 2C). Interestingly, when we compared our dataset to a recent RNA-interactome study, in which the pull-down efficiencies of phosphorylated and non-phosphorylated forms of mRNA-binding proteins were analyzed (Vieira-Vieira *et al*, 2022) (see also Materials and Methods), we found that among KIS-phosphorylated proteins there is a significant enrichment of RNA-binding proteins that decrease their binding to mRNA upon phosphorylation (Fig 2D), but not the contrary, suggesting that KIS phosphorylation would likely act by promoting dissociation of protein complexes from mRNA. In all, KIS phosphoproteomics reveals a strong association of this kinase with alternative splicing regulators, PTBP2 being one of the most relevant phosphotargets.

### KIS kinase activity counteracts PTBP2 function on exon usage

To define the role of KIS in alternative splicing in neurons, we decided to focus our study on the link between PTBP2 and KIS. It is known that loss of PTBP2 causes extensive changes in alternative splicing in brain (Vuong *et al*, 2016b). To assess whether these changes may be modified by KIS expression, we performed an enrichment analysis of PTBP2-regulated exons in our cortical samples to obtain population-wide data. We found that PTBP2-inhibited exons are significantly (FDR=0.001) enriched in KIS knockdown neurons, supporting the notion that KIS acts on AS, at least in part, by inhibiting PTBP2 activity. This analysis was also carried out with PTBP2-activated exons, but no significant enrichment was observed in this case (Fig 3A). As expected, randomization of our original RNA-seq dataset resulted in a total loss of enrichment (Fig EV3A). In addition, applying our exon usage analysis to RNA-seq data from PTBP2 KO cells (Vuong *et al*, 2016b) produced high enrichment scores for both PTBP2-inhibited and activated exons as expected (Fig EV3B), which supports our approach to analyzing exon inclusion from RNA-seq data. To validate the effect of KIS on PTBP2 activity, we analyzed the alternative splicing of CamKIIβ transcript in KIS knockdown samples (shCtrl vs shKIS). This transcript, which has important neuronal functions, undergoes exclusion of exons 13 and 16 in a PTBP2-dependent manner (Licatalosi *et al*, 2012; Li *et al*, 2014). Notably, the KIS knockdown (shKIS) increased the exclusion of these exons about 4-fold (Fig 3B). Since the observed changes could be due to changes in the expression of PTBP2 or in one of its interactors, we analyzed the expression levels of PTBP2, MATR3 and hnRNP M. No significant differences were observed between cortical neurons infected with shCtrl *vs* shKIS (Fig EV3C).

**Figure 3.**
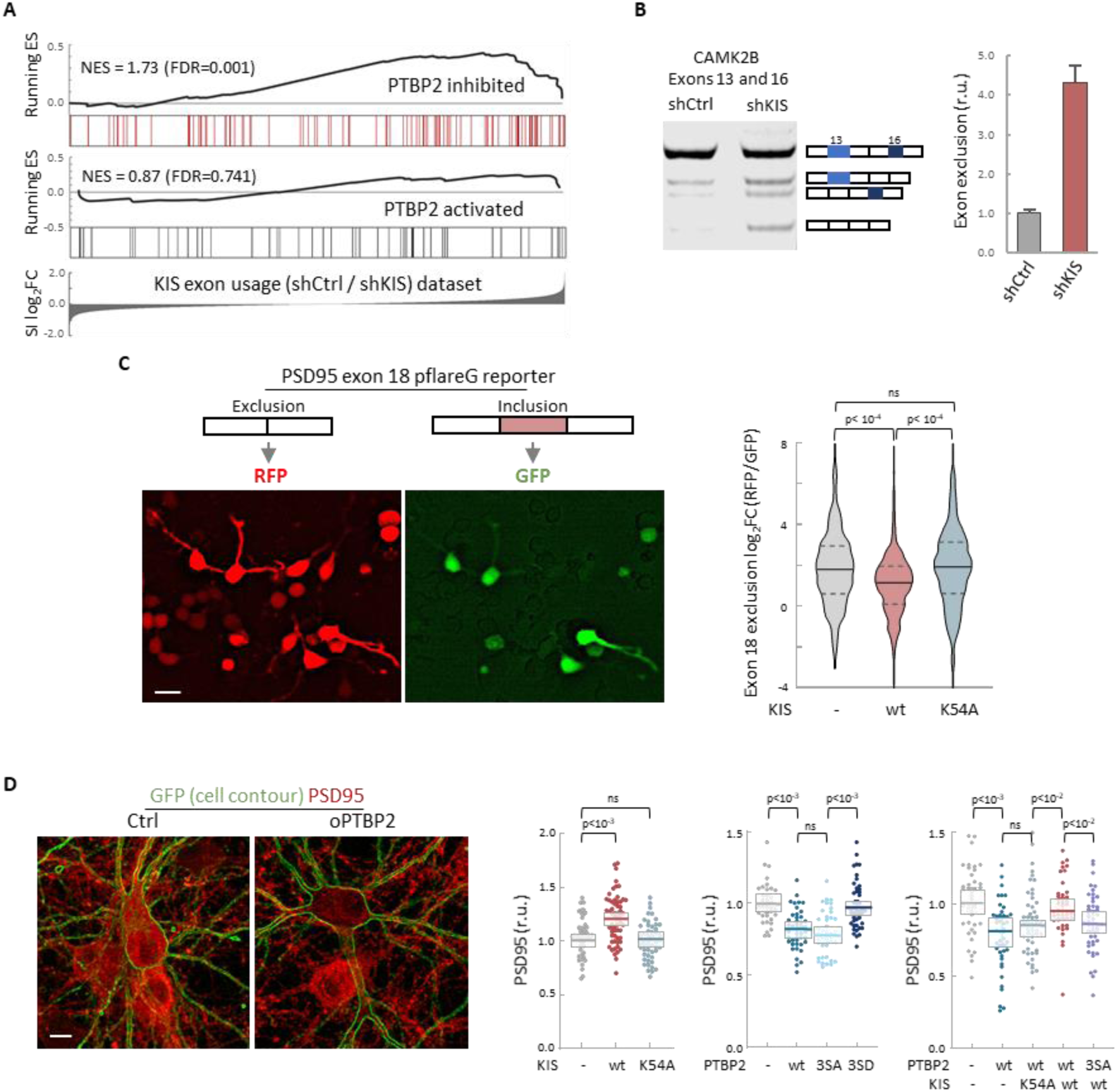
KIS kinase activity inhibits Ptbp2-exon-dependent exclusion. **A** Gene enrichment analysis of PTBP2-regulated exons. Barcode plots show the position of exons excluded (red) or included (gray) by PTBP2 within the transcriptomic dataset of exons upregulated by KIS. Normalized enrichment scores (NES) and the corresponding FDR values are shown. **B** Semi-quantitative RT-PCR of CamKIIβ from control (shCtrl) and KIS knockdown neurons (shKIS). The level of isoforms displaying exon exclusion relative to the full-length isoform is plotted as mean ± SEM (n = 3). **C** Neuroblastoma N2A cells co-transfected with dual-fluorescent reporter (Zheng *et al,* 2013) and KIS or KIS^K54A^. Exon 18 exclusion is plotted as a function of the RFP to GFP fold change (log2FC) from ctrl (n = 672), KIS (n= 1054), or KIS^K54A^ (n= 970) expressing cells. Median ± Q values and the results of a Kruskal-Wallis test are also shown. Scale bare, 20µm. **D** PSD95 immunofluorescence of hippocampal neurons transfected with vectors expressing PTBP2 and GFP as reporter. Plots show the quantification of PSD95 endogenous levels in single cells transfected with the indicated wild type and mutant proteins in three sets of experiments. Median ± Q values (n>40) and the results of a Kruskal-Wallis test are also shown. Scale bare, 10µm.

To further confirm the effects of KIS on PTBP2 activity, we used a fluorescent reporter that monitors *in vivo* alternative splicing in cultured neuroblastoma N2A cells. The minigene reporter pflareG-exon 18 of the *Dlg4* gene produces RFP and GFP from the exon-skipping and exon-including isoforms, respectively (Zheng *et al*, 2013). PTBP2 induces the exclusion of exon 18, which control *Dlg4* expression during neuronal development (Zheng *et al*, 2012). We found that KIS expression significantly decreased exon 18 exclusion levels compared to non-transfected cells or cells transfected with the K54A kinase dead mutant (Fig 3C). These results suggest that KIS inhibits PTBP2 activity on exon 18.

The *Dlg4* gene encodes PSD95, a key regulator of synaptic assembly, and the exclusion of exon 18 produces a change in the isoform expression pattern and induces mRNA degradation by NMD (Zheng *et al*, 2012). Thus, we decided to analyze the effects of KIS on PSD95 endogenous levels in hippocampal neurons. In agreement with the above experiments using the splicing fluorescent reporter, KIS overexpression produced an increase in endogenous PSD95 levels compared to control neurons or neurons transfected with the K54A mutant (Fig 3D). In addition, whereas transfection of wt or a phospho-null triple mutant (3SA) of PTBP2 produced a reduction in PSD95 levels when compared to control neurons, expression of a phosphomimetic mutant (3SD) had no significant effect (Fig 3D). Together, these results suggest that the phosphorylation state of PTBP2 strongly modulates its role on PSD95 expression. To test the effects of KIS on PTBP2 in this scenario, we co-transfected phosphorylation variants of KIS and PTBP2. First, KIS rescued the negative effects of PTBP2 on PSD95 expression levels. Second, this rescue was dependent on the kinase activity of KIS since co-transfection of PTBP2 with the inactive KIS^K54A^ mutant did not increase PSD95 expression. Finally, KIS did not rescue PSD95 levels when co-transfected with the phospho-null 3SA mutant of PTBP2 (Fig 3D). Taken together, our findings provide substantial evidence that KIS inhibits PTBP2 activity by phosphorylation.

### Phosphorylation by KIS dissociates PTBP2 protein complexes and impairs RNA binding

To further examine how phosphorylation by KIS modifies PTBP2 splicing activity, we decided to investigate whether the inhibition of PTBP2 produced by KIS could be due to alterations of protein-protein interactions within PTBP2 complexes. It is well known that PTBPs proteins are able to bind to different proteins acting synergistically (Fig 2B) and, in some cases, modify its specificity for some mRNAs (Coelho *et al*, 2016; Attig *et al*, 2018) .On the other hand, in the present study we have shown that key PTBP2 interactors are also KIS phosphotargets. Therefore, it is reasonable to speculate that the association between PTBP2 and its co-regulators could be modulated as a result of phosphorylation by KIS. To assess interactions *in vivo,* HEK293T cells were co-transfected with FLAG-PTBP2 and HA-KIS or KIS^K54A^, and endogenous levels of MATR3 and hnRNPM proteins were analyzed in FLAG immunoprecipitates. Co-transfection with KIS resulted in a significant decrease in PTBP2 binding to MATR3 and hnRNPM. In contrast, this decrease did not occur when the K54A mutant was co-transfected, indicating that the kinase activity of KIS is required to cause dissociation of protein complexes (Fig 4A). Similar results were obtained in binding assays *in vitro*, in which purified KIS forms were added to FLAG-PTBP2 immunoprecipitates, evidencing a direct participation of KIS in the dissociation of PTBP2-MATR3-hnRNPM complexes (Fig EV4A). Next, we wanted to confirm these results using an orthogonal approach, by measuring Förster resonance energy transfer (FRET) between mGFP-PTBP2 and mScarlet-hnRNPM in the nucleus of hippocampal neurons transfected at 6 DIV, when KIS expression has not yet reached maximal levels (Fig 1B). In agreement with the data from immunoprecipitation experiments, KIS transfected neurons showed a significant reduction in FRET levels compared to the KIS^K54A^ mutant (Fig 4B). Similar results were obtained for both hnRNPM and MATR3 when FRET was analyzed in the nuclei of HEK293T transfected cells (Fig EV4B). Thus, the two orthogonal approaches show that phosphorylation by KIS disturbs protein-protein interactions within splicing regulatory complexes.

**Figure 4.**
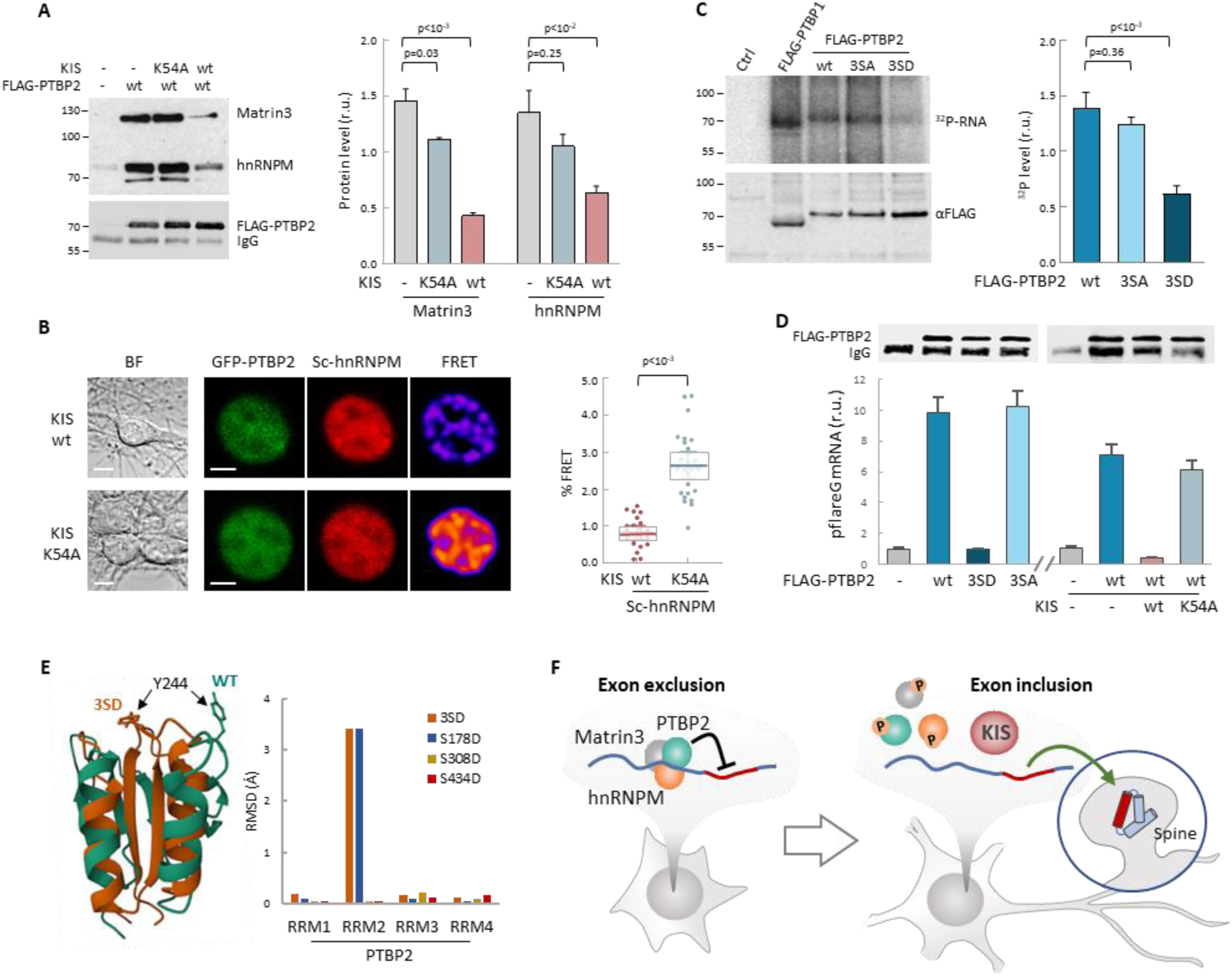
Phosphorylation by KIS disrupts PTBP2 protein complexes and compromises PTBP2 RNA binding ability. **A** Constructs expressing FLAG-PTBP2 and KIS, KIS^K54A^ were transiently transfected into HEK293T cells, and after 24 hours, cell lysates were subject to FLAG immunoprecipitation. Immunoprecipitates were analyzed by immunoblotting to detect endogenous Matrin3 and hnRNPM proteins. A representative blot is shown with empty vector as control (−). Plots show the quantification of endogenous protein levels relative to FLAG-PTBP2 in immunoprecipitates as mean ± SEM (n = 3) values. **B** Bright-field (BF), fluorescence and FRET images of representative nuclei from hippocampal neurons co-expressing GFP-PTBP2 and mScarlet-hnNRPM in combination with KIS or KIS^K54A^ proteins (scale bar, 5µm). BF images are also shown (scale bar, 10 µm). Plot shows FRET levels from single neuron nuclei expressing KIS (n = 22) or KIS^K54A^ (n = 26) as median ± Q values. The results of a Mann-Whitney test are also shown. **C** PTBP1 and PTBP2 CLIPs were performed in HEK293T cells expressing FLAG-PTBP1 and FLAG-PTBPs proteins. UV RNA crosslinked and co-precipitated with FLAG-PTBP1 and FLAG-PTBP2s was labeled with 32P-ATP. A representative radiogram gel is shown in the upper image, while the FLAG immunoprecipitates blot is shown in the bottom image. Plot shows the quantification of 32P signal relative to FLAG-PTBP2 immunoprecipitates. Bars represent mean ± SEM of n = 3 experiments. **D** PTBP2 CLIP was carried out in HEK293T cells transfected with the pFlareG-PSD95 reporter in combination with plasmid vectors expressing the indicated PTBP2 and KIS proteins. UV-crosslinked pFlareG mRNA co-precipitated with FLAG-PTBP2 was analyzed by RT-qPCR and normalized relative to FLAG-PTBP2 in immunoprecipitates (upper blots). Bars represent mean ± SEM (n = 3). **E** AlphaFold2 prediction of the RRM2 domain from wild-type (green) and 3SD (orange) PTBP2. The different orientation of Tyr244 is highlighted. Plot shows the root-mean-squared deviation (RMSD) values of the four RRM domains in the different phosphomimetic PTBP2 mutants. **F** PTBP2 protein complexes inhibit gene expression through exon exclusion in neuronal precursors whereas, in differentiated neurons, phosphorylation by KIS dissociates PTBP2 protein complexes, which, by being released from the pre-mRNA, allow inclusion of the exon. The contraposition of both mechanisms could be essential to fine-tune expression patterns in differentiated neurons with diversified synaptic connections.

Finally, we wanted to test whether phosphorylation by KIS could also affect the RNA binding capacity of PTBP2. For this purpose, we applied the cross-linking and immunoprecipitation CLIP method to HEK293 cells transfected with FLAG-PTBP2 wt or phosphorylation mutants. Notably, we immunoprecipitated a much lower amount of endogenous RNA crosslinked to the phospho-mimetic mutant 3SD compared to FLAG-PTBP2 wt or 3SA mutant proteins (Fig 4C). Next, we used the same CLIP approach with HEK293 cells transfected with the pflareG-exon 18 reporter (as in Fig 3D), and analyzed levels of the reporter mRNA in PTBP2 immunoprecipitates. In agreement with the previous CLIP experiment, the reporter mRNA levels were strongly reduced in phospho-mimetic 3SD pulldown samples. Moreover, we observed that the reporter mRNA levels in FLAG-PTBP2 immunoprecipitates were reduced when KIS was co-expressed, whereas the KIS^K54A^ mutant had no effect (Fig 4D). These results reinforce the notion that phosphorylation of linker regions within PTBP2 may affect the interactions between RRM domains and RNA. Interestingly, AlphaFold2 predicts important structural alterations in the phosphomimetic S178D and triple 3SD mutants particularly affecting RRM2, which has been shown to function as the minimal splicing repressor domain of PTBP2 (Robinson & Smith, 2006) (Fig 4E and Fig EV4C).

## Discussion

Alternative splicing has emerged as a critical and widespread gene regulatory mechanism across organisms and tissues. Among the broad tissue types present in metazoans, the nervous system contains some of the highest levels of alternative splicing. Alternative splicing controls multiple steps of neuronal development and plasticity and is a causal agent of pathology (Furlanis & Scheiffele, 2018). We had previously shown that KIS is required for the proper expression of postsynaptic proteins and the correct development of synaptic spines (Cambray *et al*, 2009; Pedraza *et al*, 2014), suggesting that KIS could be involved in the mechanisms that generate protein diversity by alternative splicing. Although many documented examples of alternative splicing regulation exist, much remains to be understood about the control of specific splicing regulators originating tissue and neuron subtype-specific splicing patterns (Raj & Blencowe, 2015). In the present study, we show that phosphorylation by KIS counteracts the activity of the splicing regulator PTBP2 and, as a consequence, controls gene isoform expression during neuronal differentiation and connectivity (Fig 4F).

Splicing site recognition and pre-mRNA splicing are dynamic processes involving constant rearrangement of ribonuclear proteins on the pre-mRNA being processed. Splicing regulatory proteins contain RNA-binding and protein-protein interaction domains. They bind with low specificity to mostly single-stranded parts of the pre-mRNA. To overcome the low RNA binding specificity, splicing regulatory proteins use their protein interaction domains to bind to each other. Most splicing factors are post-translationally modified, phosphorylation being an important contributor to the splicing code that governs the fate of a pre-mRNA. Phosphorylation may induce a global conformational change in a protein that may allosterically either promote or inhibit protein−protein or protein−RNA interactions. Among the best studied classes of splicing proteins regulated by phosphorylation are the SR proteins (serine/arginine rich splicing factors) (Stamm, 2008). For example, phosphorylation of the SR protein SF2/ASF increases its binding to U1 snRNP and decreases its binding to the RNA export factor TAP/NXF1. Moreover, the ability of its arginine/serine domain (RS) to bind to RNA also depends on its phosphorylation state(Huang *et al*, 2004). The RS domain is highly disordered and switches to a compact, partly ordered arched shape upon phosphorylation (Thapar, 2015). Bioinformatics studies have shown that phosphorylation of intrinsically disordered regions (IDRs) occurs more frequently than in folded domains. This could be due to their increased flexibility and therefore improved kinase accessibility. In accordance with this idea, recombinant PTBP1 and PTBP2 under *in vitro* splicing conditions present different phosphate modifications located in the unstructured N-terminal and linker regions (Pina *et al*, 2018). Although the underlying structural mechanism is not clear, the presence of a phosphoryl group introduces electrostatic charges that can affect the structural stability of binding regions or their ability to interact with RNA or other proteins (Newcombe *et al*, 2022). Supporting this idea, the current study shows that the phosphorylation of PTBP2 linker regions by KIS dissociates both protein-protein and protein-RNA interactions in the PTBP2 complex. Interestingly, AlphaFold2 predicts that a mutation mimicking Ser178 phosphorylation, located at the end of linker 1, would produce major structural changes in the RRM2 domain (Fig 4E and Fig EV4C). This domain is of particular relevance for the interaction with RNA and splicing co-regulators, such as Matrin3, Raver1 or hnRNPM (Coelho *et al*, 2015). Within the RRM2 domain, Tyr244 is particularly critical for these interactions (Joshi *et al*, 2011). The AlphaFold2 prediction shows that a phosphomimetic mutation in Ser178 completely changes the spatial orientation of this Tyr244. AlphaFold2 is not yet reliable enough to predict structures with posttranslational modifications, such as glycosylation, methylation and phosphorylation (Bertoline *et al*, 2023). Thus, we cannot rule out that phosphorylation of other residues (Ser308 and Ser434) does not cause important structural alterations in PTBP2. In particular, AlphaFold2 fails when the modifications lie within unstructured regions, as is the case of Ser308 (Chakravarty & Porter, 2022).

It is worth pointing out that Ser178 and Ser434 are also present in PTPB1, suggesting that phosphorylation of PTBP1 by KIS may have a role in tumorigenesis. In gastric cancer, KIS up-regulation promotes growth and migration, indicating that KIS functions as an oncogene (Feng *et al*, 2020). In hepatocellular cancer, KIS stimulates the expression of specific cell cycle regulation genes. Furthermore, the KIS interactome in liver cancer cells revealed a significant enrichment of potential KIS interactors in mRNA splicing, including hnRNPM and PTBP1 (Wei *et al*, 2019). Finally, KIS has been associated with the epithelial-mesenchyme transition by phosphoproteomic and RNA-seq analyses (Arfelli *et al*, 2023). Thus, in a non-neuronal context, it is plausible that phosphorylation by KIS modulates the capacity of PTBP1 to interact with RNA and co-regulatory proteins to control cell proliferation. One of the new PTBP1 interactors is SUGP2 (SURP and G-patch domain containing 2), which shares high homology with SUGP1. SUGP2 has been shown to interact with the RRM2 region of PTBP1 in HeLa nuclear extracts (Coelho *et al*, 2015, 2016). It is worth mentioning that SUGP1 is one of the most enriched proteins in our KIS phosphoproteome (see Fig 2A). It forms part of the spliceosome complex, interacts with the general splicing factor U2AF2 and has been reported to play an important role in branch recognition by its association with SF3B1. Altered levels of SUGP1 have been found in a large number of uveal melanoma and breast cancer cells (Zhang *et al*, 2019). In the future it would be interesting to explore whether phosphorylation by KIS interferes with SUGP-PTBP interactions and the functional relevance of this mechanism in cancer and neurogenesis.

The PTBP family represents one of the most salient regulators of splicing in neurogenesis. Sequential changes in the expression of PTBP1 and PTBP2 allow for the establishment of distinct splicing regulatory patterns during neuronal maturation (Zheng, 2016; Fisher & Feng, 2022). PTBP2 expression decreases during the first postnatal week of brain development, maintaining a moderate level in adulthood (Li *et al*, 2007; Fisher & Feng, 2022). Accordingly, our RNA-seq analysis showed significant PTBP2 expression in cortical neurons at 18 DIV (Fig EV2A). Strikingly, PTBP1 knockdown stimulates PTBP2 expression and is sufficient to induce neuronal differentiation of fibroblasts (Xue *et al*, 2013). On the other hand, PTBP2 depletion in developing mouse cortex causes degeneration of this tissue over the first three postnatal weeks, a time when the normal cortex expands and develops mature circuits. Cultured Ptbp2−/− neurons fail to mature and die after a week in culture (Li *et al*, 2014). However, despite all these data on the physiological relevance of PTBP2 in neurogenesis, the role of this splicing regulator in differentiated neurons and how its activity is regulated are still not understood. Regarding neuronal differentiation, it is worth noting that KIS expression has been shown to increase during brain development (Bièche *et al*, 2003), coinciding with postnatal decrease of PTBP2 levels (Zheng *et al*, 2012). Therefore, it is reasonable to speculate that during neuronal differentiation, in addition to transcriptional downregulation, phosphorylation by KIS contributes to harnessing PTBP2 activity. In mature neurons, alternative splicing has a well-established role in expanding proteome diversity (Mauger & Scheiffele, 2017). In this regard, the contraposition of two mechanisms in splicing may represent a fine-tuning mechanism to modulate proteome diversity as a function of plasticity-inducing signals. In this regard, single-cell transcriptomic data from hippocampal neurons show that expression variability of KIS and PTBP2 is much higher compared to actin (Perez *et al*, 2021) (Fig EV4D). Differences in the expression of these two splicing regulators at a single neuron level would contribute to increasing diversity in neural circuits as a crucial property in information processing (Miller *et al*, 2019).

## Materials and Methods

### Primary dissociated cultures

Animal experimental procedures were approved by the ethics committee of the National Research Council of Spain (CSIC). Hippocampi and cortex were dissected from E17.5 CD1 mice (undetermined sex) in HBSS containing 0.6% glucose and 10 mM HEPES. After dissection, tissues were trypsinized (Fisher, 15090046) in HBSS at 37°C for 15 min. Enzymatic digestion was stopped by washing the tissue three times with MEM (Fisher, 31095029) supplemented with 10% FBS and 0.6% glucose. Hippocampi were left to sediment between washes, and centrifugation was avoided to keep cell viability. Trypsin-treated tissue was then mechanically disaggregated by passing through a flame-polished Pasteur pipette (∼10 times). Cells were plated at desired density on Poly-D-lysine (Merck-Sigma, P7886)-coated plates (0.5 mg/ml Poly-L-lysine in borate buffer, pH 8.5) and maintained in MEM, 10% FBS and 0.6% glucose for 2–4 h. Culture medium was then substituted for Neurobasal (Fisher, 11570556) supplemented with 2% B27 (Fisher, 11530536) and 1% GlutaMAX (Fisher, 35050-038). Primary hippocampal cultures were transfected using CalPhos mammalian transfection kit (Clontech, 631312) as previously (Jiang & Chen, 2006).

### Cell culture and transfection

HEK293T cells and neuroblastoma N2A cells were cultured in DMEM medium with 10% FBS. HEK293T transfection was carried out with Lipofectamine 2000 (Invitrogen, 11668030) and N2A transfection was performed with FuGene HD (Promega, E2311), according to the manufacturer’s instructions.

### DNA constructs

Site-directed mutagenesis in PTBP2 and KIS cDNAs was performed by In-Fusion HD (Takara, 638909). pcDNA3-FLAG, pEGFP-C3, mScarlet-C1, pGEX-KG, pET28a and pcDNA3-HA were used as host vectors. Plasmids were prepared using NucleoSpin Plasmid Kit (Macherey-Nagel, 740588) for cell line transfections and NucleoBond Xtra Midi Plus EF Kit (Macherey-Nagel, 740422) for neuron transfections.

### Lentivirus production and infection

The shRNA sequence in pLKO.3-GFP lentiviral vector against mouse KIS was GAGTGCGGAGAATGAGTGTTT (MISSION shRNA library, TRCN0000027622) and control non-mammalian shRNA was from Merck-Sigma (SHC002). HEK293FT cells were transfected with Lipofectamine 2000 with lentiviral vector, envelope plasmid pVSV-G, and packaging plasmid pHR’82ΔR, and cultured in DMEM 10% FBS. Lentiviruses were harvested 1, 2 and 3 days after transfection, filtered through 0.45 μm cellulose-acetate syringe filters, and concentrated by centrifugation at 26,000 rpm for 2 h at 4°C. Titration was performed by serial dilution infection in HEK293T and checking the transduction efficiency by fluorescence as described(Ritter *et al*, 2017). Cortical neurons cultured for 7DIV were infected with 2 MOI lentiviruses for 3h, and harvested at 18 DIV.

### RT-PCR and semiquantitative PCR and qPCR

RNAs were extracted from indicated samples using EZNA total RNA purification KIT (Omega, R6834-01), digested with RNase-free DNase (Roche, 11119915001), and 1 ug of RNA was reverse-transcribed into cDNAs using the Prime Script RT Reagent Kit (Takara, RR037B) according to the manufacturer’s protocol. RT-PCR primers designed to amplify two spliced isoforms with different sizes are shown in Table EV1. was performed with Taqman probes (6xFAM-BQ1) on LightCycler@96 Real-Time PCR system (Roche) according to manufacturer’s instructions. qPCR primers are shown in Table EV1.

### Co-Immunoprecipitation, in vitro phosphorylation, and western blot

HEK293T cells where transfected using Lipofectamine 2000 with a proportion 1:3:9 of pEGFP, pcDNA-FLAG/FLAG-PTBP2 and pcNBM470-HA-KIS/HA-KIS^K54A^. After 48 hours post-transfection, cells were wash with cold phosphate buffered saline (PBS) and homogenized with lysis buffer (50mM Tris HCl pH 8, 150mM KCl, 5mM MgCl2, 0.25% NP-40, 0.05% sodium deoxycholate, 5% glycerol, 1mM EGTA, 1mM EDTA, and protease and phosphatase inhibitor). To preserve only direct interactions, RNA was digested using RNase I (Fisher, 10330065). After sonication, lysates were clarified by centrifugation at 12000g and incubated 2h at 4°C with pre-equilibrated α-FLAG M2 affinity gel beads (Merck-Sigma, A2220). Beads were washed three times with lysis buffer. After washes, beads were eluted for western blot analysis or equilibrated to perform in vitro phosphorylation assay. For in vitro assay, FLAG-immunoprecipitates were incubated in BK buffer (50mM HEPES pH 7.6, 10mM MgCl2, 2mM DTT, 10µM ATP) with 200 ng of His-KIS or His-KIS^K54A^ purified from *E. coli* for 30min at 30°C. After three washes with lysis buffer, beads were eluded with SDS sample buffer. Antibodies used for immunoblotting were: α-FLAG M2 (1:500; Merck-Sigma, F3165), α-Matr3 (1:500; BioNova, A300-591A-T), α-hnRNP M3/4 (1:500; BioNova, A303-910A-T), IRDye 680RD (1:10000; LI-COR Biosciences, 926-68070) and α-rabbit HRP (1:10000; Fisher, 31460, RRID AB_228341).

### In vitro kinase assay

Kinase reactions were carried out in 20 μl of BK buffer (50mM HEPES pH 7.6, 10mM MgCl2, 2mM DTT, 10µM ATP), 10 μCi of [γ-32P]-ATP, 0.2 μg of GST-KIS (or GST-KIS^K54A^), and 2 μg of the corresponding GST-PTBP2 proteins. Kinase reactions were incubated for 20 min at 30°C. Phosphorylated products were separated by SDS-PAGE, stained with Coomassie Brilliant Blue, dried, and analysed by autoradiography.

### Immunofluorescence

Hippocampal neurons were fixed at different DIVs using 4% PFA and 4% sucrose in PBS for 30 min at 4°C and then washed three times with PBS. Neurons were permeabilized for 5 min with 0.1% Triton X-100 and blocked with 5% NGS in PBS (blocking solution). Primary antibody α-PSD95 (EMD Millipore, MABN68, RRID: AB_10807979) were diluted 1:400 in blocking solution. Proteins were detected by incubation with secondary antibodies Alexa 568 donkey anti-mouse (Life Technologies, A10037, RRID: AB_11180865) diluted 1:1000 in blocking solution. Images were acquired with a Zeiss LSM780 confocal microscope using the following parameters: ∼15-μm-thick stacks were imaged every 0.5 μm (pinhole set at 1 Airy unit), under 40× objective (0.13 μm/pixel). Laser power and PMT values were kept constant throughout images and conditions. PSD95 somatic quantification was performed using ImageJ by measuring fluorescent intensity on transfected neurons and relativizing it to the somatic expression of the non-transfected neurons in the same field(Zheng *et al*, 2012). For nuclear SRRM2 immunofluorescence, HEK293T were transfected with GFP or GFP-KIS plasmids. 24 hours post-transfection, cells were fixed with 4% paraformaldehyde for 20 min. Plates were washed twice with PBS and permeabilized with cold 100% methanol for 2 minutes, before sequential staining with primary and secondary antibodies. Cells were washed three times with PBS and imaged within 24h. Primary immunofluorescence was staining with α-SRRM2 (1:250; Merck-Sigma, HPA066181) and secondary with Alexa 568 α-rabbit (1:1000; Life Technologies, A10042; RRID: AB_2534017).

### Dual fluorescence labelling

N2A cells were reverse-transfected with a proportion 3:9 of pFlareG-PSD95-e18(Zheng *et al*, 2013) and HA-KIS or HA-KIS^K54A^. After 72h fluorescent levels were recorded with a Leica Thunder Imager. Led power and exposition time were kept constant for all images and conditions. After image-deconvolution process, single cell RFP and GFP were measured with ImageJ. The ratio of exclusion/inclusion of exon 18 in pFlareG was measured as the log2(RFP/GFP).

### PTBP1 and PTBP2 CLIP assay

HEK293T cells where transfected with FLAG-PTBPs proteins. 48h post transfection, cells were washed once with PBS and crosslinked with 150mJ/cm2 UV light at 254nm in Stratalinker 2400. Cells where collected by centrifugation and resuspended on lysis buffer (50mM Tris-HCl ph7.4, 100mM NaCl, 1% IGEPAL CA-630, 0.1% SDS, 0.5% sodium deoxycholate, protease and phosphatase inhibitors). After sonication, lysates where digested with 4U/mL Turbo DNase (Fisher, AM2238) and 1.5U/μL RNase I (Fisher, 10330065). Lysates were centrifugated and supernatants incubated with pre-equilibrated α-FLAG M2 affinity gel beads for 2h at 4°C. Beads were washed twice with high-salt wash buffer (50 mM Tris–HCl, pH 7.4, 1 M NaCl, 1 mM EDTA, 1% IGEPAL CA-630, 0.1% SDS, 0.5% sodium deoxycholate, protease and phosphatase inhibitors). At that point, beads were equilibrated twice with PNK buffer (20 mM Tris–HCl, pH 7.4, 10 mM MgCl2, 0.2% Tween-20) and RNA 5’ ends where labelled with [γ-32P]-ATP and T4 PNK (NEB, M0201L) at 37°C for 5 min. Beads were eluded at 70°C for 5 min with NuPAGE loading buffer (Fisher, 11549166) and loaded in 4–12% NuPAGE Bis-Tris gel (Fisher, 10247002) for subsequent electrophoresis and transference to nitrocellulose membrane (Merck-Sigma, GE10600003). Precipitated RNA was analyzed by autoradiography and FLAG-PTBP protein levels were measured by western blot.

### pFlareG-PSD95 CLIP assay

Protocol used for CLIP assay from pFlareG plasmid was essentially as in the previous section with some modifications. Briefly, HEK293T cells were transfected with pFlareG-PSD95 reporter in combination with plasmid vectors expressing FLAG-PTBP2 proteins and HA-KIS proteins. After crosslinking, cells where resuspended in lysis buffer plus 40U/ml RNase inhibitor (Attendbio, RNI-G), sonicated and incubated with α-FLAG M2 affinity gel beads overnight at 4 °C. Beads were washed twice with lysis buffer, twice with lysis buffer without detergents and once with DNase buffer (10Mm Tris-HCl, pH 7.4, 2,5 Mm MgCl2, 0.5 Mm CaCl2). Beads were digested with DNase I (Merck-Sigma, DN25) for 15 min and inactivated. RNAs bound to the beads were used as a template for cDNA synthesis using Maxima H Minus cDNA first strand synthesis Kit (ThermoFisher, 10338179) and random hexamers as primers. 2µl of cDNA was used as a template in a real-time qPCR assay. pFlareG mRNA was amplified with GFP annealing primers (see Table EV1). Relative levels of pFlareG were calculated for each condition using GAPDH as a housekeeping gene and normalized relative to FLAG-PTBP2 protein levels immunoprecipitated checked by western blot using α-FLAG M2 antibody (1:500; Merck-Sigma, F3165) and IRDye 800RD (1:10000; LI-COR Biosciences, 926-32210).

### FRET imaging

Hippocampal cultures were transfected at 6 DIV with FRET biosensor plasmids expressing mGFP-PTBP2 proteins and pmScarlet-hnRNPM, together with HA-KIS or HA-KIS^K54A^, as indicated. Live imaging was conducted 16 hours after transfection. Neurons were live-imaged using a Zeiss LSM780 confocal microscope with a 40× 1.2-NA water-immersion objective. Images were 1024 × 1024 pixels, with a pixel width of 65 nm. Briefly, donor (mGFP-PTBP2) protein was excited at 488 nm, and its emission was measured at 490 to 532 nm (ID). Excitation of the acceptor (pmScarlet-hnRNPM) was at 561 nm, and emission was measured at 563 to 695 nm (IA). We also measured the total signal emitted at 563 to 695 nm when excited at 488 nm (IF) to obtain the FRET efficiency as F% = 100 * (IF − kD*ID − kA*IA) / IA, kD, and kA, correcting acceptor cross-excitation and donor bleed-through, respectively, with the aid of FRETmapJ, a plugin that also provides maps with the FRET signal as pixel value for local quantification. For HEK293T experiments, cells were transfected with plasmids expressing mGFP-PTBP2 and pmScarlet-hnRNPM or pmScarlet-Matr3, together with HA-KIS or HA-KIS^K54A^, as indicated.

### Mass spectrometry-based phosphoproteomics analysis

HEK293T cells were transfected with plasmids expressing KIS and KIS^K54A^. Methods used for sample preparation and data acquisition were as previously described(Azman *et al*, 2023) with some modifications. Briefly, 100 µg of total lysate were used for filter aided sample preparation (iFASP). Digestion with trypsin and chymotrypsin was performed in two different experiments before TMT isobaric labelling (TMT-126TM and TMT-131TM Thermo Fisher). Trypsin (Promega, V5280) and chymotrypsin (Promega, V106A) digested eluates were then pooled independently. The two pooled mixtures were fractionated into four fractions using PierceTM High pH reverse-phase fractionation kit (Life Technologies, 84868), according to the manufacturer’s instructions. Fractions samples were subjected to TiO phosphopeptide enrichment using TiO enrichment kit (GL Sciences, 5010-21308), according to the manufacturer’s instructions. Phosphopeptides were analysed using a Q Exactive plus Orbitrap mass spectrometer (Barts Cancer Institute, London). MaxQuant (version 1.6.3.3) software was used for database search and label-free quantification of mass spectrometry raw files. The search was performed against a FASTA file of Homo sapiens proteome, extracted from Uniprot (2016). All downstream data analysis was performed using Perseus (version 1.6.2.3). To perform enrichment analysis, using GSEA software (Mootha *et al*, 2003), phospho-site information was summarized at protein level by phosphoprotein score (PPS). PPS was calculated as the sum of the log2FC value for each phospho-site on a given protein. For Gene Ontology analysis, the c5.all.v2023.1.Hs.symbols.gmt list provided by GSEA was utilized. Top 50 PTBP2 interactors were obtained from the STRING database by full string network.

### RNA extraction, sequencing and data analysis

Total RNA was obtained from 18 DIV infected cortical neurons and 7 and 18 DIV of untreated neurons using EZNA total RNA purification kit (Omega, R6834-01), and used to prepare polyA+ enriched libraries for paired-end (2x100 bp) sequencing (BGI DNBSEQ platform). More than 90 million reads were obtained per sample, which were aligned to the mm10 mouse genome using HISAT2 (Kim *et al*, 2015) in the Galaxy platform (Boekel *et al*, 2015). Assignment to transcriptional units, standard filtering steps and basic statistical analysis of triplicate samples was done using Rsubread (version 3.6) (Liao *et al*, 2019) in Bioconductor (Huber *et al*, 2015) and custom R scripts. Genes with a single exon were removed and only exons with more than 5 reads in all samples were considered. For every annotated exon a splicing index (SI) (Mauger *et al*, 2016) was calculated as the ratio of exon to gene rpkm. GO term and specific gene set enrichment analysis was performed using Enrichr (Chen et al, 2013) and GSEA (Mootha *et al*, 2003) tools, respectively. A subset of 8457 exons with low variability (Fano factor<0.0005) within the KIS knockdown triplicate samples was selected, and the significance of differences to control triplicate samples was assessed by calculating the false-discovery rate (FDR) with a modified t-test statistic that incorporates a background variance parameter s0=0.1 (Goss Tusher *et al*). In this manner, a total of 2247 exons offered q<0.05. In contrast, when the same analysis was applied to the experimental dataset randomized by Gaussian attribution or permutation, less than 5 exons displayed q<0.05.

### Statistical analyses

Statistical analyses were performed using GraphPad PRISM (version 9). Multiple comparisons were performed with a Kruskal-Wallis test, and the resulting pairwise p values are shown in the corresponding figure panels. Data are displayed as median and quartile (Q) values, the number of samples is described in the figure legends. Protein levels by immunoblotting and mRNA levels by RT-PCR were determined in triplicate samples, and mean ± SEM values are shown.

### Software

FRETmapJ can be obtained as ImageJ (Wayne Rasband, NIH) plugin from https://www.ibmb.csic.es/en/department-of-cells-and-tissues/control-of-local-mrna-expression/

### Data and materials availability

The mass spectrometry proteomics data have been deposited to the ProteomeXchange Consortium through the PRIDE partner repository with the dataset identifier (pending). RNA-seq fastq files are available as BioProject (pending). Plasmids (Table EV1) not deposited in Addgene are available upon request.

## Acknowledgments

We thank E. Rebollo for technical assistance, M. Aldea for the helpful comments, and C. Rose for editing the manuscript. We also thank members of M. Aldea and C. Gallego laboratories for the stimulating discussions and ideas. Funding: This work was funded by a grant from the Ministry of Science and Innovation of Spain and the European Union (FEDER) (PID2020-113231GB-I00) to C.G. Medical Research Council UK (MR/P009417/1) and Barts Charity (MGU0346) grants to FKM.

## Author contributions

**M.M.A**., **R.O.** and **C.G** conceived and designed the study; **M.M.A**., **M.B.M**, **A.M.N**, **R.O**., and **C.G** performed bench experiments; **M. D.** and **F.K.M.** contributed to phosphoproteomics analysis; **M.M.A.** and **C.G.** analyzed the data; **C.G.** supervised the project, acquired the funding and wrote the manuscript.

---

**Figure EV1.**
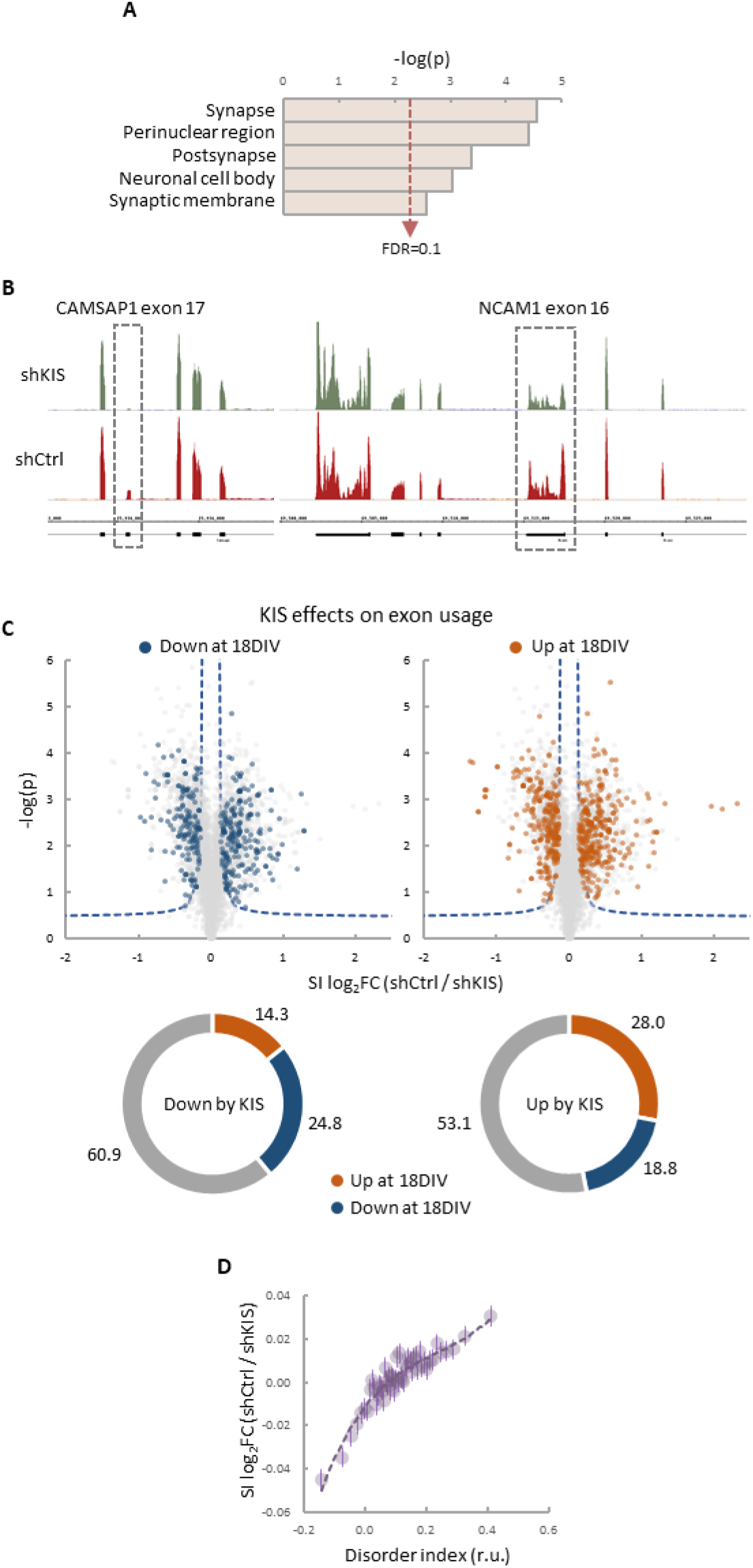
KIS promotes exon inclusion in genes encoding proteins related to synaptogenesis and neurogenesis. **A** GO term enrichment analysis of genes with more than one exon upregulated by KIS. FDR value is shown. **B** Representative read profiles showing KIS-dependent exons in CAMSAP1 and NCAM1 genes (dashed-line rectangles). **C** Volcano plots show t-test significance (-log2p) as a function of splicing index (SI) fold change (log2FC) in control neurons (shCtrl) compared to KIS knockdown neurons (shKIS). Significantly (FDR < 0.05) downregulated (blue) and upregulated (orange) exons during neuronal differentiation are highlighted. Sunburst charts show the percentage of exons differentially modulated during differentiation in sets downregulated (left) and upregulated (right) by KIS. **D** Exon usage fold change (log2FC) in control neurons (shCtrl) compared to KIS knockdown neurons (shKIS) as a function of peptide disorder index (see Materials and Methods). Mean values of binned data (bin size = 2500 exons, purple circles) with confidence limits for the mean (α = 0.05) and a regression line are shown.

**Figure EV2.**
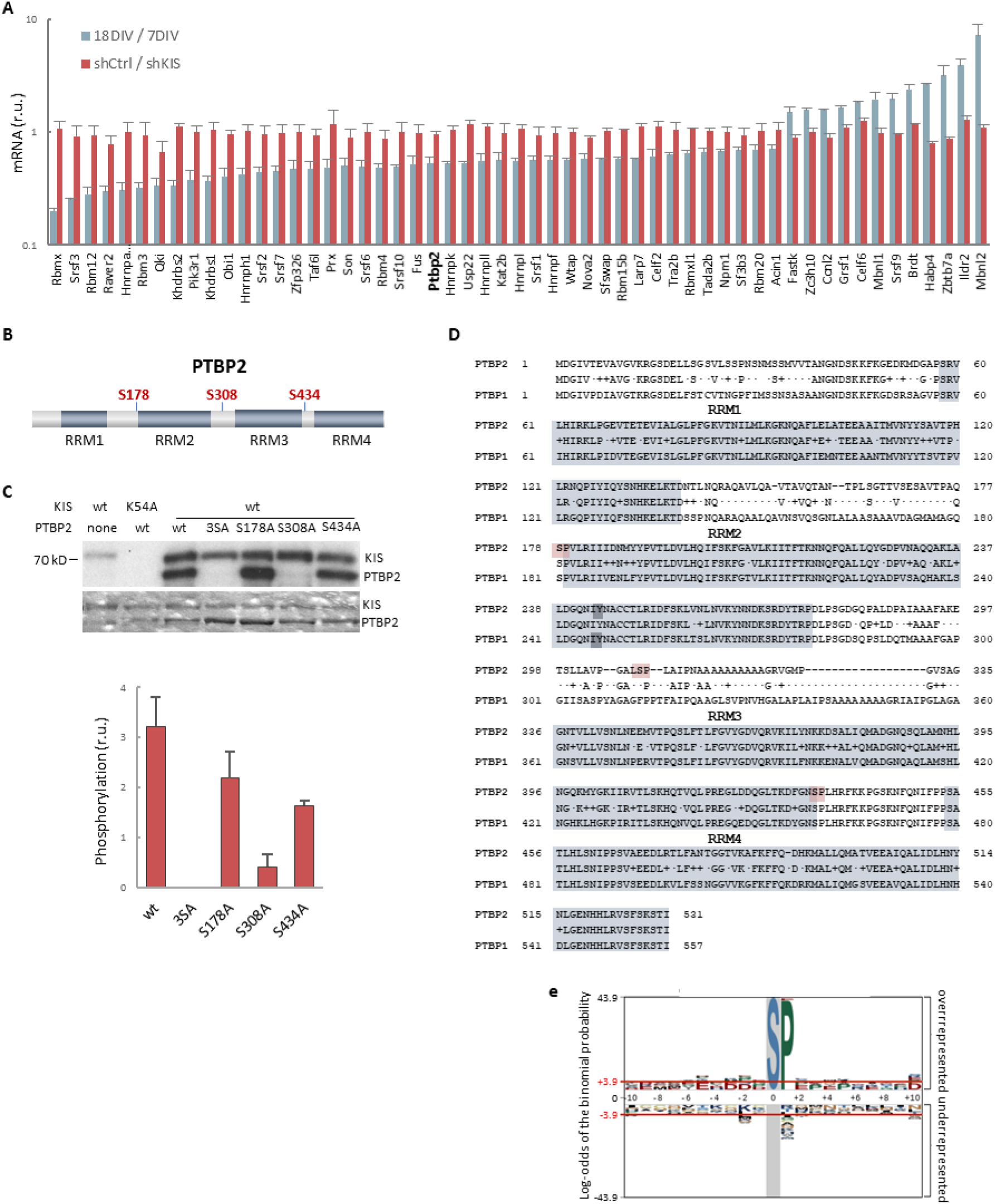
In vitro phosphorylation of PTBP2 by KIS. **A** mRNA levels of splicing regulators from GO biological process in cortical neurons at 18DIV relative to 7DIV (blue) or expressing shKIS relative to shCtrl (red). Mean ± SEM (n = 3) values from genes with log2FC >0.5 and FDR < 0.1 are shown. **B** PTBP2 protein scheme showing RNA recognition motifs (RRMs) and localization of KIS phosphorylated serines within linker regions. **C** KIS in vitro kinase assay of indicated GST recombinant proteins purified from *E.coli*. Representative autoradiography (top) and Coomassie staining (bottom) images are shown. The plot shows the quantification of phosphorylation signals as mean ± SEM (n = 3) values. **D** Aligned amino acid sequences of mouse PTBP2 and PTBP1. RNA recognition motifs (RRMs) are highlighted in gray and KIS phosphorylated serines in red. Tyr244 in RRM2 is highlighted in bold type. **E** pLogo analysis (O’Shea *et al,* 2013). KIS preferentially phosphorylates serines (S) with high frequency (89.38%) followed by proline (P) at position +1 with a frequency of 41.67% and statistical significance (p < 0.05). Input sequences = 143.

**Figure EV3.**
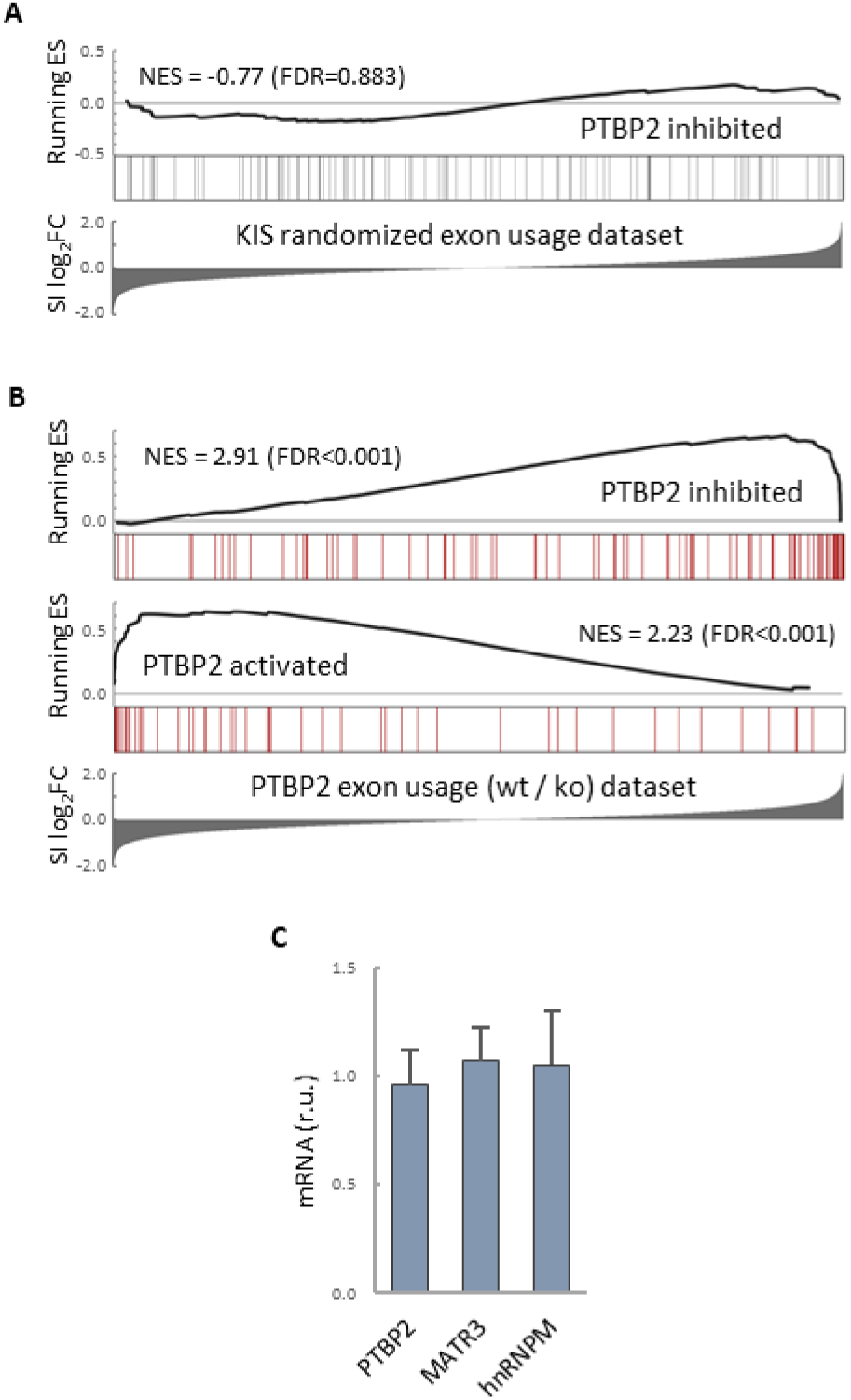
PTBP2 RNA-seq analysis. **A** Gene enrichment analysis of PTBP2-regulated exons. Barcode plots show the position of exons excluded by PTBP2 within a KIS randomized exon usage dataset. Normalized enrichment scores (NES) and the corresponding FDR values are shown. **B** Gene enrichment analysis of PTBP2-regulated exons applying our exon usage analysis (see Materials and Methods). Barcode plots show the position of exons excluded and included by PTBP2 within the transcriptomic dataset from PTBP2 KO cells (Vuong *et al*, 2016b). Normalized enrichment scores (NES) and corresponding FDR values are indicated. **C** Quantification of mRNA levels from KIS knockdown cortical neurons relative to control neurons. Bars represent mean ± SEM (n = 3).

**Figure EV4.**
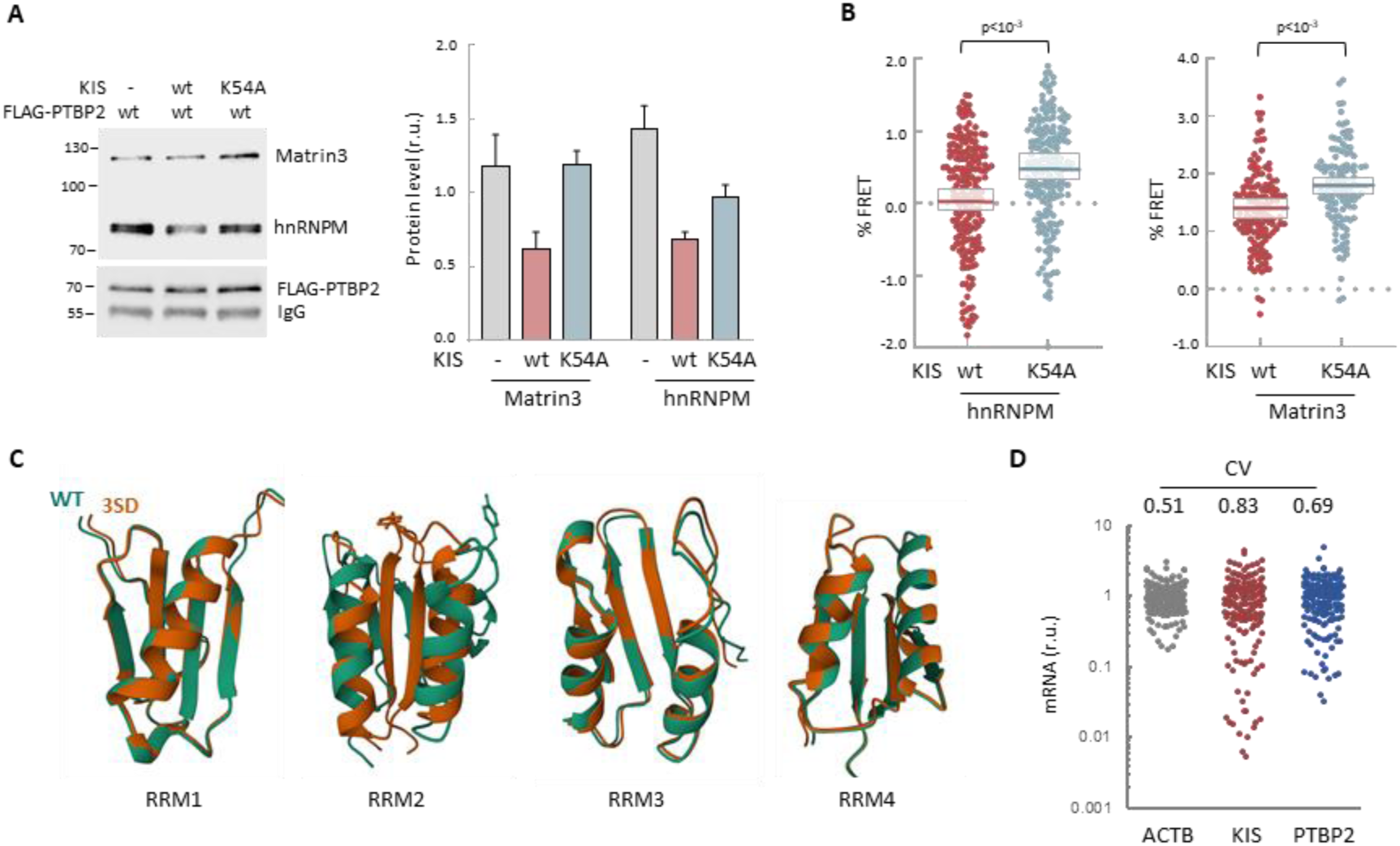
In vitro and in vivo experiments showing dissociation of splicing complexes by KIS kinase activity. **A** Cell lysates from HEK293T cells expressing FLAG-PTBP2 protein were incubated with KIS recombinant proteins purified from bacteria as His fusions. FLAG immunoprecipitates were analyzed by immunoblotting to detect endogenous Matrin3 and hnRNPM proteins. A representative blot is shown with empty vector as control (−). Plots show the quantification of endogenous protein levels relative to FLAG-PTBP2 in immunoprecipitates as mean ± SEM (n = 3). **B** HEK293T cells co-expressing GFP-PTBP2 and mScarlet-hnNRPM or mScarlet-Matrin3 in combination with HA-KIS or KIS^K54A^ proteins. Plots show the FRET levels from single neuron nuclei as median ± Q values (n>100). The results of a Mann-Whitney test are also shown. **C** AlphaFold2 prediction of the four RRM domains from wild-type (green) and 3SD (orange) PTBP2. **D** Expression variability of KIS and PTBP2 in single-cell transcriptomic data from hippocampal neurons (Perez *et al,* 2021). The plot shows relative levels of actin, KIS and PTBPT2 mRNAs from single cell somas. The coefficient of variation (CV) is indicated at the top.

